# Evolutionary dynamics of RNA viruses during persistent infection in Drosophila melanogaster

**DOI:** 10.64898/2026.11.11.765870

**Authors:** Anamarija Butković, Hervé Blanc, Mauro Castelló-Sanjuán, Maria-Carla Saleh, Rubén González

**Affiliations:** Institut Pasteur, Université Paris Cité, Viruses and RNA Interference Unit (Paris, France)

## Abstract

Persistent viral infections provide an extended window for viral evolution, yet their intra-host trajectories across the host lifespan remain underexplored. Here, we tracked the longitudinal evolution of three single-stranded RNA (ssRNA) viruses (Drosophila A virus [DAV], Drosophila C virus [DCV], and Nora virus) and one double-stranded RNA (dsRNA) virus (Bloomfield virus) during persistent infection in *Drosophila melanogaster*. By analyzing viral populations across early, intermediate, and late infection stages, we mapped dynamic shifts in the viral quasispecies and the emergence of adaptive mutations. These data demonstrate that long-term persistence drives viral evolution in a viral species-specific manner. DAV evolution was defined by clonal interference, driven by the continuous turnover of competing haplotypes. Nora virus exhibited prolonged evolutionary stasis punctuated by a late-stage sweep, suggesting adaptation to the senescing host environment. In contrast, standard DCV populations were locked under strong purifying selection, where no adaptive mutations were detected. However, this constraint collapsed in high-titer DCV infections, triggering extensive mutational diversification. Distinct from the ssRNA dynamics, the multi-segmented dsRNA Bloomfield virus maintained higher sequence fidelity and a stable genomic formula. Ultimately, these results demonstrate that viral evolution during persistent infection is governed by the interplay between specific host-pathogen interactions and the physiological decline of the host.

## INTRODUCTION

Unlike acute infections that quickly resolve through host clearance or death, persistent infections maintain active viral replication over extended periods. This chronicity provides an expanded arena for mutational exploration, allowing variants to continuously arise, compete, and potentially fix within the population (Vignuzzi et al. 2006, Holmes 2009, Domingo 2020, Farci et al. 2000). Hence, persistent infections may favor the emergence of variants with altered virulence, tissue tropism, or transmission properties (Holland et al. 1979, Frielle et al. 1984, Domingo 2020, Bonneaud and Longdon 2020). While the evolutionary capacity of RNA viruses across serial passages in invertebrate hosts is well established (Mongelli et al. 2022, Lezcano et al. 2023, Castiglioni et al. 2024), how intra-host dynamics unfold longitudinally within a single persistently infected, aging host remains largely unexplored.

The fruit fly *Drosophila melanogaster* provides a highly tractable system to dissect these longitudinal evolutionary dynamics (Dionne and Schneider, 2008). Wild and laboratory *Drosophila* populations harbor diverse viral communities that can establish stable, long-term persistent infections (Goic and Saleh 2012, Webster et al. 2016, Obbard et al 2020, Wallace et al. 2021). These infections drive diverse physiological costs: reducing host lifespan, accelerating the host aging, altering reproductive output, disrupting the community of microbes associated with the host, and modulating behavior (Wallace and Obbard 2024, Castelló-Sanjuán et al. 2025, González et al. 2026a).

*Drosophila* viruses belong to diverse taxonomic families and exhibit distinct infection strategies, providing valuable models for studying the spectrum of virus-host evolutionary dynamics (Webster et al., 2015). In this study, we focus on four enteric RNA viruses: Drosophila A virus (DAV, family *Permutotetraviridae*, single genomic segment of ∼4.8 kb), Drosophila C virus (DCV, family *Dicistroviridae*, single genomic segment of ∼9.2 kb), Nora virus (family *Noraviridae*, single genomic segment of ∼12.2 kb), and Bloomfield virus (family *Reoviridae*, segmented genome with multiple segments). DAV, DCV, and Nora are positive-sense, single-stranded RNA viruses, while Bloomfield virus is a double-stranded RNA virus. To isolate their individual impacts, we previously established controlled Drosophila monoinfection lines. Utilizing these lines, we demonstrated that although the four viruses share a common oral-fecal transmission route, they differ substantially in the infection responses they induce and in their pathogenicity: DAV and DCV are highly pathogenic, whereas Nora and Bloomfield viruses exhibit milder pathogenicity (Castelló-Sanjuán et al., 2025). This system provides a uniquely controlled environment to characterize viral evolutionary dynamics over the course of a single host’s adult lifespan.

Here, we leverage a longitudinal RNA-seq dataset (Castelló-Sanjuán et al., 2025) to track the evolutionary trajectories of these four viruses across the lifespan of persistently infected *D. melanogaster* adults. Because these viruses were maintained within the same fly genotype (*w^1118^*) for several generations prior to being sequenced, they represent viral populations adapted to their host. By analyzing viral populations across early, intermediate, and late infection stages, we map temporal fluctuations in nucleotide diversity, identify genomic regions under selection, and document the emergence of novel mutations and defective viral genomes (DVGs).

## RESULTS

### Viral genetic diversity is shaped by species-specific genomic and temporal dynamics

Single-stranded RNA (ssRNA) viruses undergo error-prone replication that fuels rapid genetic diversification (Lauring and Andino 2010). To determine how this baseline property is shaped by intra-host persistence, we quantified nucleotide diversity (π) across the genomes of DAV, DCV, and Nora virus at early (1 day post-eclosion, dpe), intermediate (12 dpe), and late (50% host mortality: 25 dpe for DAV, 28 dpe for DCV, and 52 dpe for Nora virus) infection stages (Figure 1).

**Fig 1.**
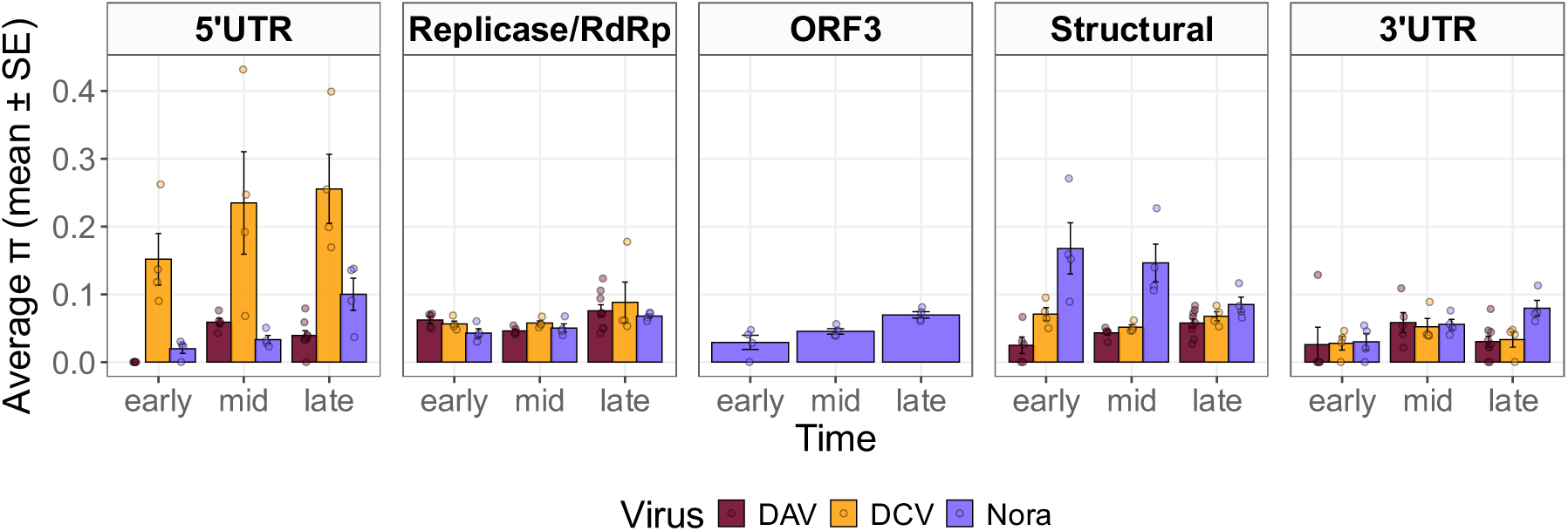
Viral genetic diversity during persistent infection. Nucleotide diversity (π) was measured across genomic regions for Drosophila A virus (DAV, red), Drosophila C virus (DCV, yellow), and Nora virus (purple) at early (1 dpe), intermediate (12 dpe), and late (median time to death) stages of infection. Each panel corresponds to a specific genomic region. Bars represent the mean nucleotide diversity ± standard error (SE), and points show the values from individual biological replicates.

Within the untranslated regions (UTRs), the 5’UTR was a major hotspot for diversity in DCV, where π increased throughout the infection. In contrast, DAV and Nora virus maintained consistently low 5’UTR diversity. Across all three viruses, the 3’UTR remained a region of low and relatively stable diversity.

In the coding regions, two distinct patterns emerged. The viral replicase (RdRp) and main structural protein coding regions of DAV and DCV maintained low, stable nucleotide diversity across all time points. Nora virus displayed a contrasting spatial profile: its main structural protein region harbored the highest baseline diversity across the genome, remaining elevated throughout the infection. The Nora virus ORF3, which encodes the capsid-stabilizing protein VP3, exhibited a steady increase in nucleotide diversity from early to late infection stages.

### Longitudinal allele frequency dynamics of the viral quasispecies

The longitudinal progression of a persistent infection subjects the viral population to a dynamic selective landscape. Because these shifting pressures drive the continuous accumulation and purging of genetic variants, we tracked the intra-host mutational trajectories of each virus by quantifying allele frequency changes (ΔAF) over time (Figure 2, Supplementary File 1).

**Fig 2.**
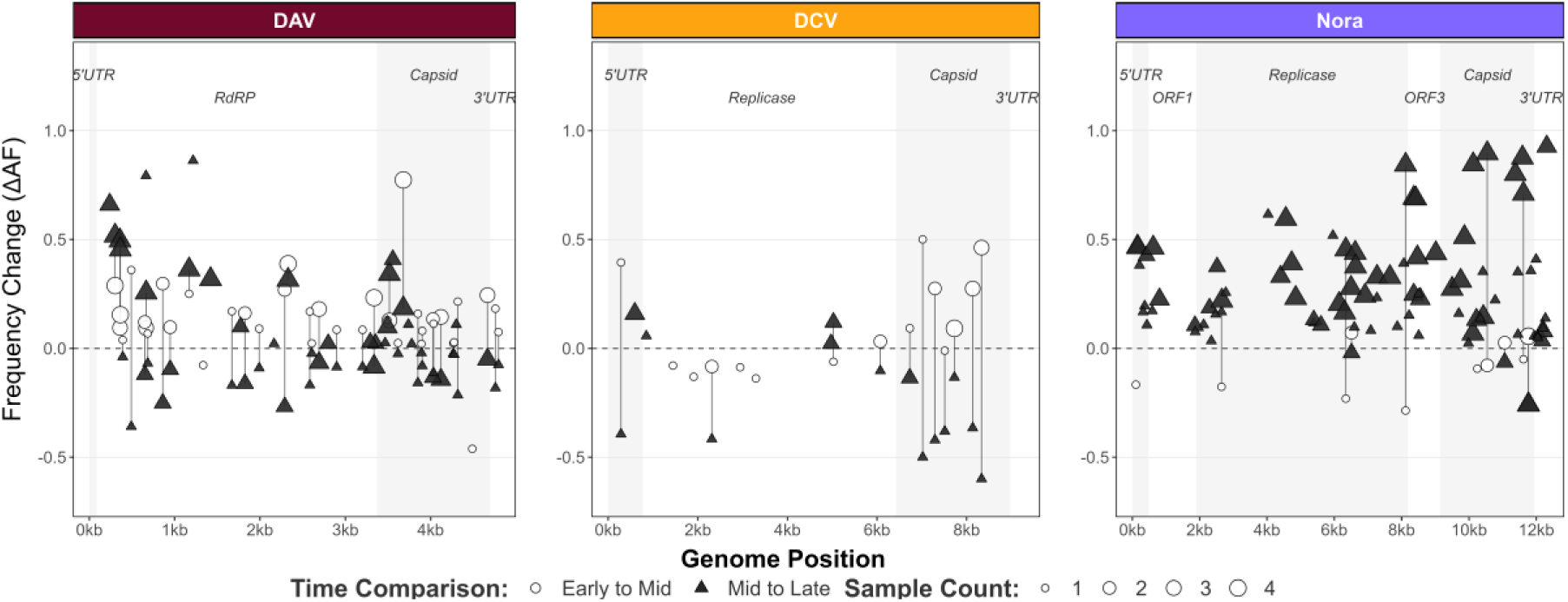
Distinct evolutionary trajectories of viral mutations during persistent infection. Plots show the change in allele frequency (ΔAF) of mutations along the genomes of DAV, DCV, and Nora virus. Open white circles represent frequency changes from the early to the intermediate time point, while filled black triangles represent changes from the intermediate to the late time point. The size of each point is proportional to the number of replicate samples (1-4) in which the mutation was detected. Dashed line at y=0 indicates no change in frequency.

The overall mutational load differed among viruses. DAV displayed minimal early mutational activity (0.5 ± 0.5 mutations at early), followed by a stable load between the intermediate (17.8 ± 4.0 mutations) and late (16.8 ± 1.9 mutations) infection stages. DCV showed a constrained pattern with lower, decreasing loads across infection (6.8 ± 0.9 at early, 4.3 ± 2.8 at intermediate, and 3.7 ± 0.3 mutations at late stages). Nora virus began with minimal mutational activity (4.2 ± 1.1 at early and 3.0 ± 0.4 mutations at intermediate stages) before undergoing rapid diversification by the late stage, expanding to 50.5 ± 3.4 mutations. Analysis of allele frequency changes revealed distinct directional trajectories. In DAV, the dynamics shifted from being dominated by mutations increasing in frequency early on (38 of 40) to a state of balanced frequency changes later (23 increasing vs. 29 decreasing). DCV’s trajectory shifted from balanced dynamics (8 increasing vs. 7 decreasing) to a reduction in high-frequency variants at the late stage (4 increasing vs. 10 decreasing). Nora virus shifted from a slight reduction in variants (4 increasing vs. 7 decreasing) to a landscape overwhelmingly dominated by frequency increases, with 81 mutations rising versus only 4 decreasing at the late stage. Complete fixation of single mutations did not occur in standard-load DAV, DCV, or Nora virus populations across these time points.

### Adaptive viral mutations emerge over the course of persistent infection

Within a heterogeneous quasispecies, positive selection acts on individual variants that enhance viral replication or stability within the host. To identify the specific genetic drivers of intra-host adaptation, we quantified the temporal trajectories and fitness effects of individual mutations across the viral genomes. This approach identifies variants conferring significant positive fitness advantages. Consistent with a landscape dominated by strong purifying selection, no novel adaptive mutations were detected in standard-titer DCV populations. In contrast, both DAV and Nora virus acquired multiple significant variants (Supplementary File 2).

In DAV, five distinct non-synonymous variants showed significant fitness advantages (Figure 3A) and increased in frequency across all infection stages (Figure 3B). To determine how these variants evolved relative to one another, we performed genetic linkage mapping (Supplementary Figure 1). The analysis revealed positive and negative correlations among the mutations, reflecting clonal interference, where distinct sub-lineages continuously compete for dominance without reaching fixation. Mapping of these mutations revealed changes within key functional domains (Figure 3D). Note that mutations are designated by their original nucleotide, genomic position, and substituted nucleotide, with the resulting amino acid alterations indicated in parentheses for non-synonymous changes. In the DAV RNA-dependent RNA polymerase (RdRp), three high-fitness non-synonymous mutations: the C239T transition (P51L), the C302T transition (T72I, driving a polar-to-hydrophobic shift), and the C364T transition (R93C), localized directly to the three functional loops of the fingers domain. Thermodynamic stability profiling via ThermoMPNN quantified changes in folding free energy (ΔΔG), indicating that these amino acid changes exert minimal destabilizing effects on the core tertiary structure, with values clustering near the neutral baseline across the replicase and capsid stability heatmaps. Evaluating these substitutions within a broader biophysical framework (Supplementary Fig. 2) revealed that adaptive mutations in DAV are predominantly placed in structurally malleable regions, where even moderately disruptive changes (ΔΔG > 0) are tolerated to drive substantial gains in relative fitness (reaching up to 1.3-fold relative to WT). Because the fingers domain orchestrates template and NTP entry, substrate binding, channel architecture, and catalytic cleft closure, these amino acid changes alter dynamic molecular interfaces to directly enhance RdRp processivity. Concurrently, the DAV capsid acquired significant non-synonymous mutations localized strictly to the N-terminal region, including the A3492G transition (K42R) and the C3516A transversion (P50H, introducing a basic aromatic ring), exhibiting minimal thermodynamic disruption. As the N-terminal domain governs nucleic acid binding during capsid assembly, these targeted substitutions can potentially optimize encapsidation efficiency. The accumulation of these non-synonymous mutations exhibited a strong positive correlation with intra-host viral load (R = 0.7, p = 0.012; Figure 3C). This functional adaptation was accompanied by a set of six significant synonymous mutations (T360C, T666C, C1170T, C1422T, T2328C, and A3679G), which emerged concurrently and displayed an equally robust correlation with viral titer (R = 0.81, p = 0.002; Supplementary Figure 3).

**Fig 3.**
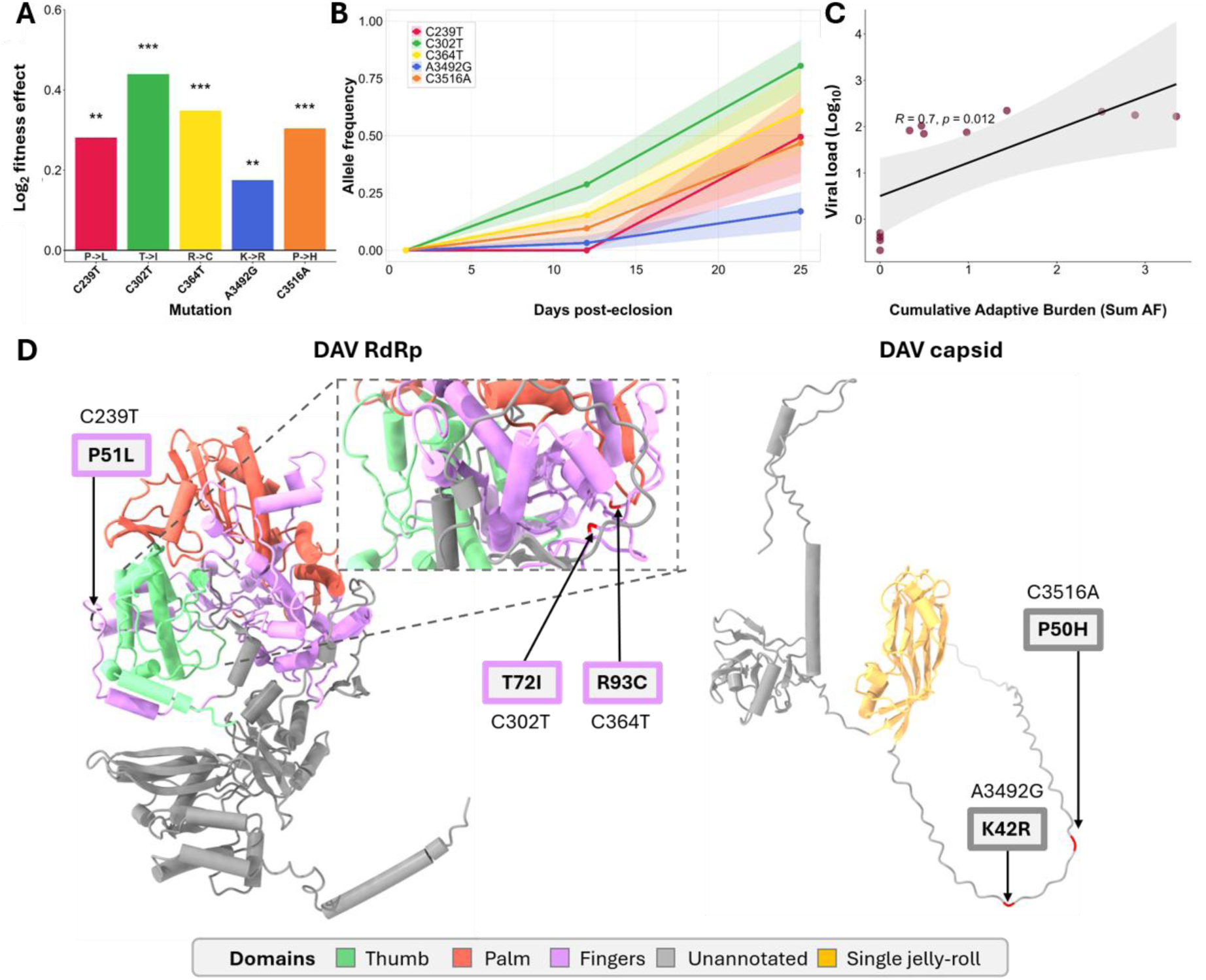
Evolutionary dynamics and fitness impacts of adaptive mutations in DAV. **(A)** Fitness effects of all 5 significant non-synonymous adaptive mutations during DAV persistence. Asterisks denote statistical significance. **(B)** Temporal allele frequency trajectories of these variants across days post-eclosion. Shaded regions represent the standard error of the mean across independent biological replicates, illustrating the simultaneous expansion of multiple competing haplotypes. **(C)** Pearson correlation between the cumulative adaptive mutational burden (sum of allele frequencies) and intra-host viral load. The solid black line indicates the linear regression fit, and the shaded area represents the 95% confidence interval (R=0.7, *p*=0.012). **(D)** Three-dimensional structural topography of the DAV RNA-dependent RNA polymerase (RdRp) and capsid proteins, with distinct structural domains colored according to the panel legend. Identified high-fitness amino acid substitutions are mapped in red to highlight specific structural regions targeted by positive selection.

Nora virus exhibited a contrasting temporal signature defined by a delayed, synchronized adaptive burst. Following evolutionary stasis during the first 12 days post-eclosion, seven non-synonymous variants exhibited significant longitudinal trajectories: five variants conferring positive fitness advantages swept the population in parallel, displacing a pre-existing minor subpopulation carrying two linked capsid substitutions (G11766A and A11767T; Figure 4A, B). Genetic linkage analysis (Supplementary Fig. 1) suggests that, rather than forming a fragmented swarm of competing lineages, these variants co-segregated as a single haplotypic block that displaced minor competing subpopulations. Structural annotation (Figure 4D) revealed that all three significant VP4 capsid trajectories localized to surface-exposed loops: the sweeping positive-fitness substitution G11617A (S825N) and the displaced subpopulation carrying two consecutive substitutions at codon 875, G11766A (E875K) and A11767T (E875V). hermodynamic profiling demonstrated that while all three substitutions exert a net stabilizing effect on the protein structure (ΔΔG < 0), they occur within distinct local structural contexts; S825N resides in a structurally malleable region where substitutions across adjacent residues induce minimal thermodynamic perturbation, whereas E875K and E875V are embedded within a structurally sensitive region flanked by residues that are highly intolerant to mutation (Supplementary Figure 2). Because the VP4 surface region is highly susceptible to host protease cleavage, this stabilization provides a direct structural mechanism for host recognition evasion. Within the Nora virus replicase, severe biochemical alterations, such as the major G4735A charge reversal (E946K) and the A6634G transition (I1579V), emerged exclusively outside the conserved core helicase and RdRp domains, exhibiting a neutral thermodynamic effect on global structure. In contrast to DAV, Nora virus adaptive mutations partitioned distinctly across biophysical categories (Supplementary Fig. 2A): while several capsid and ORF3 substitutions localized to sensitive regions where they exerted net stabilizing effects (ΔΔG < 0), others exploited malleable structural backgrounds to maximize fitness without crossing the critical unfolding threshold of 2.0 kcal/mol. Finally, the capsid-stabilizing protein ORF3 acquired two significant mutations (A8358G, N57S; and C9012T, T275I) localizing to the beta-sheet region and the long alpha-helix, both conferring enhanced structural stability to the protein. The accumulation of these non-synonymous variants exhibited a highly significant linear correlation with intra-host viral load (R = 0.94, p < 0.001; Figure 4C). This functional adaptation was accompanied by an expansive set of 29 significant synonymous and 4 non-coding mutations, which underwent an identical, synchronized late-stage sweep and displayed an equally robust correlation with viral load (R = 0.95, p < 0.001; Supplementary Figure 4).

**Fig 4.**
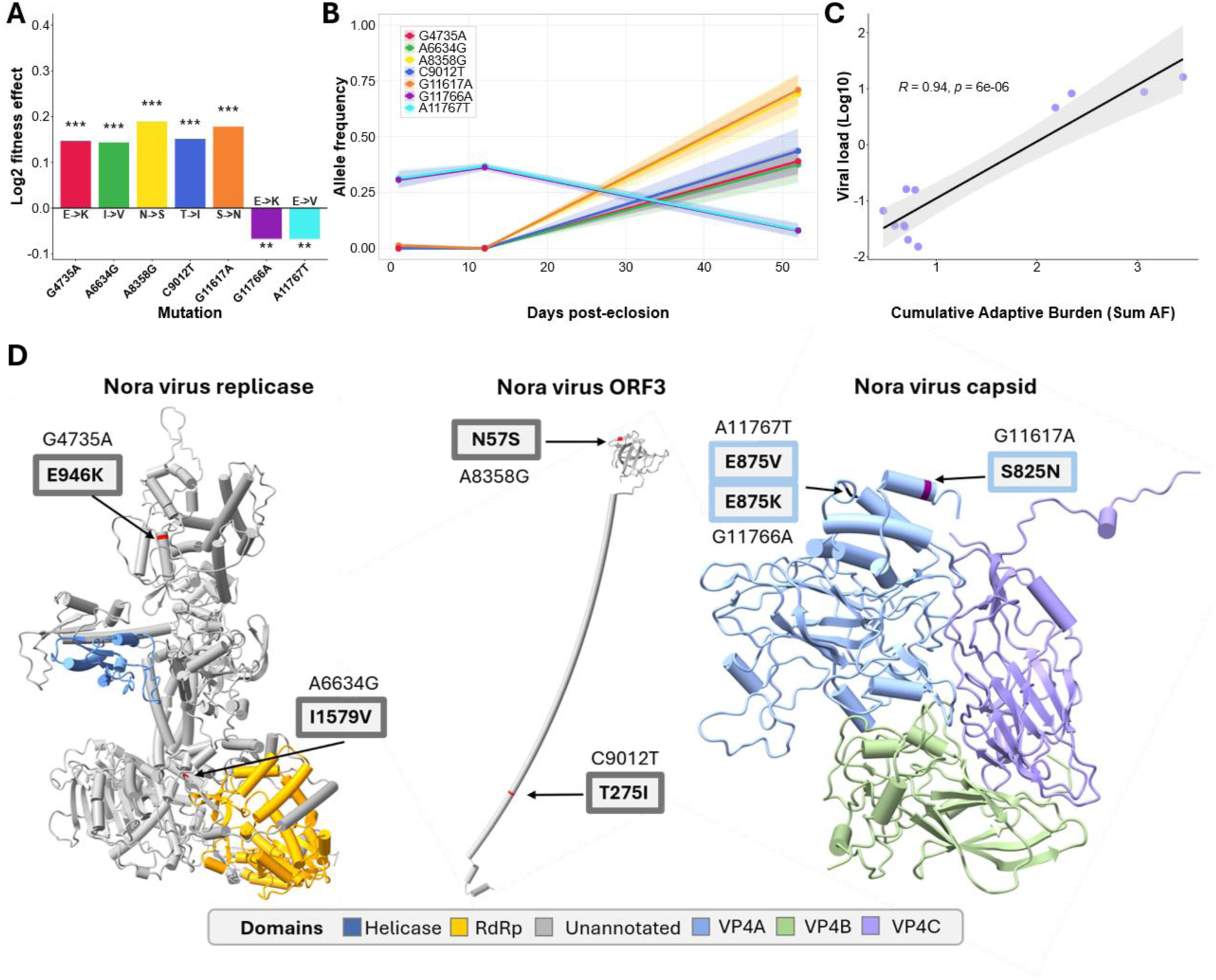
Evolutionary dynamics and fitness impacts of adaptive mutations in Nora virus. **(A)** Fitness effects of all 7 significant non-synonymous adaptive mutations emerging during Nora virus infection. Asterisks denote statistical significance. **(B)** Temporal allele frequency trajectories of these specific variants across the host lifespan (days post-eclosion). Shaded regions represent the standard error of the mean across biological replicates, capturing the delayed, synchronized selective sweep at the late infection stage. **(C)** Pearson correlation between the cumulative adaptive mutational burden (sum of allele frequencies) and intra-host viral load. The solid black line indicates the linear regression fit, and the shaded area represents the 95% confidence interval (R=0.94, *p*=6×10^-6^). **(D)** Three-dimensional structural topography of the Nora virus replicase, ORF3, and capsid proteins, with distinct structural domains colored according to the panel legend. Identified amino acid substitutions are mapped in red to highlight specific structural regions remodeled during the late-stage selective sweep.

### The magnitude of viral replication determines DCV mutational diversity

Viral evolutionary trajectories depend heavily on census population size, as higher replication rates expand the supply of *de novo* mutations. Whereas most DCV-infected flies maintain controlled viral titers, a distinct subset develops viral loads several orders of magnitude higher than baseline (Castelló-Sanjuán et al., 2025). To determine how replication magnitude alters the balance between purifying selection and mutational diversification from the onset of host mortality (6 dpe; Figure 5, Supplementary File 3), we sequenced these high-load populations and compared their longitudinal mutational landscapes against standard-load controls.

**Fig 5.**
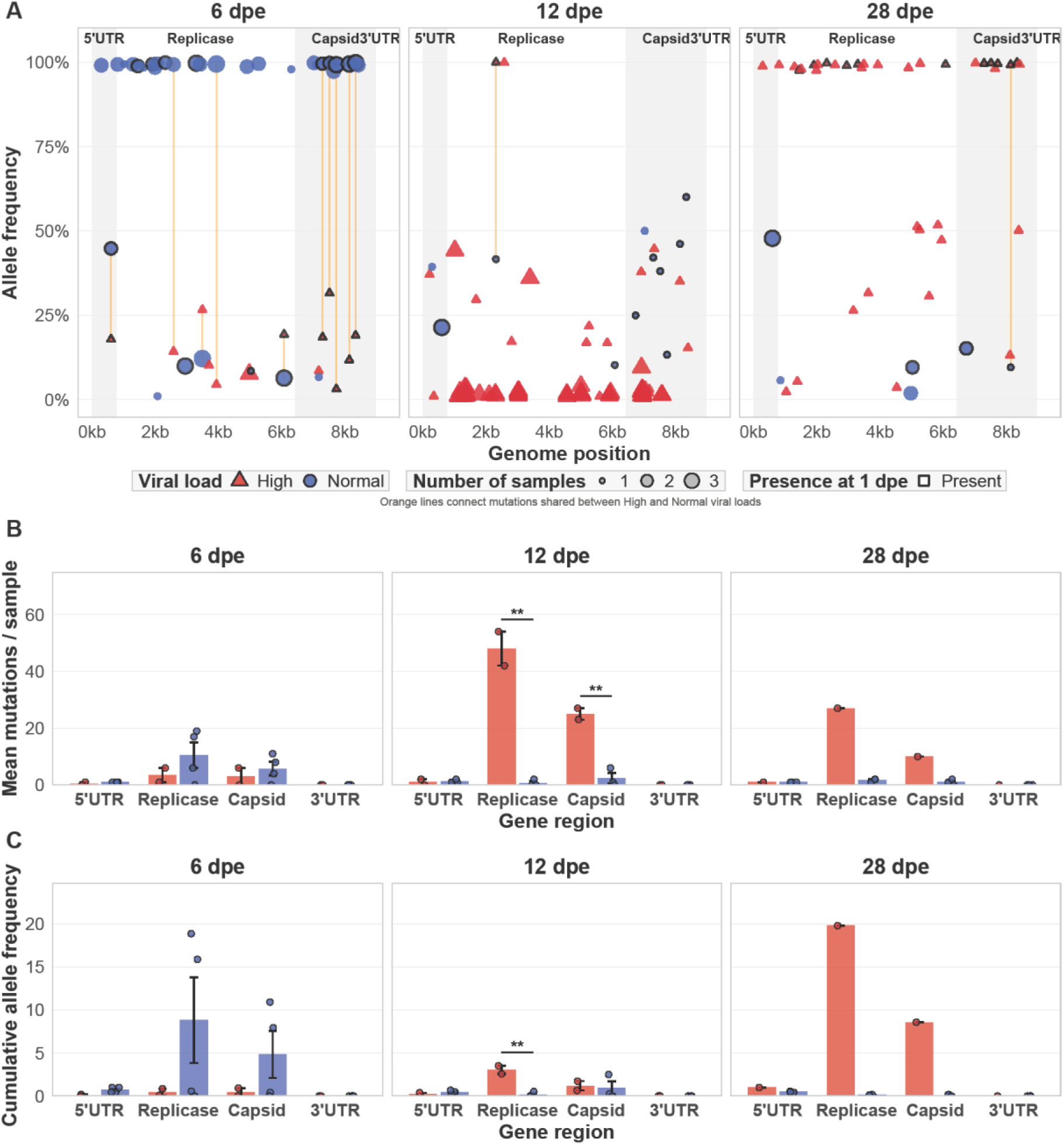
Viral load magnitude dictates the evolutionary trajectory of DCV mutations. Viral populations from standard-load (blue circles/bars) and high-load (red triangles/bars) infections were analyzed at 6, 12, and 28 dpe. (A) Allele frequencies of individual mutations mapped across the DCV genome. Point size is proportional to the number of biological replicates in which a given mutation was detected (*n* = 3-4 for standard-load; *n* = 3 at 6 dpe, *n* = 2 at 12 dpe, and *n* = 1 at 28 dpe for high-load). Black borders denote genetic variants already present in the ancestral 1 dpe population. Solid orange lines connect mutations shared across both load groups. (B) Mutational richness, represented as the mean number of discrete mutations per sample categorized by genomic region. (C) Mutational burden, quantified as the Cumulative Allele Frequency (Sum AF) per sample, capturing the overall penetrance and fixation weight of the mutant swarm. For both bar plots, error bars represent the standard error of the mean (SEM), and individual data points denote independent biological replicates. Statistical significance between standard and high-load populations within each gene region and time point was determined using a two-tailed Student’s t-test with pooled variance.

At 6 dpe, the two viral load groups exhibited distinct mutational architectures. Standard-load populations were dominated by multiple mutations at near-fixation (>90% allele frequency), whereas high-load populations maintained genetic variants at low frequencies. As the infection progressed to intermediate (12 dpe) and late (28 dpe) stages, these evolutionary trajectories completely diverged. In standard-load infections, both mutational richness (mean number of mutations) and mutational burden (cumulative allele frequency) shrank over time. Conversely, high-load populations exhibited a massive expansion in mutational diversity at 12 dpe. This spike was primarily driven by variants in the replicase and capsid genes, resulting in a significantly higher mutational richness compared to standard-load populations. By 28 dpe, in the single surviving high-load biological replicate available at this late time point, this expanded mutant swarm reached near-fixation across multiple novel loci in the replicase and capsid genes.

### Defective viral genome populations exhibit dynamic, virus-specific profiles

Defective viral genomes (DVGs) shape viral pathogenesis by interfering with wild-type replication and modulating host responses (Rezelj et al. 2021). To determine if these truncated and rearranged variants exist during persistent infection in Drosophila, we mapped DVG formation, abundance, and structural recombination across all time points (Supplementary File 4).

DVG abundance and structural profiles exhibited strict virus- and load-dependent dynamics across infection stages (Fig. 6A). For DCV, DVGs were undetected at early stages under high viral load but were detected in the standard load populations at 6 dpe (detected in 2 of 4 analyzed samples), generating 508 unique variants with breakpoints distributed broadly across the ∼9.2 kb genome alongside two prominent peaks. At the intermediate stage (12 dpe), DCV populations expanded to 1,860 variants (detected in both high-load samples) (Fig. 6A). Progression into the high-load late stage (28 dpe, single available sample) revealed continued persistence with extensive structural clouds off the main diagonal. Conversely, DVG formation in DAV was restricted to 2 of 4 replicates at the late stage (25 dpe), yielding 9 unique variants at basal read depths. DAV DVGs localized exclusively to the RdRp gene without the broad structural dispersion observed in DCV. For Nora virus, DVGs emerged exclusively at the late infection stage (52 dpe).

**Fig 6.**
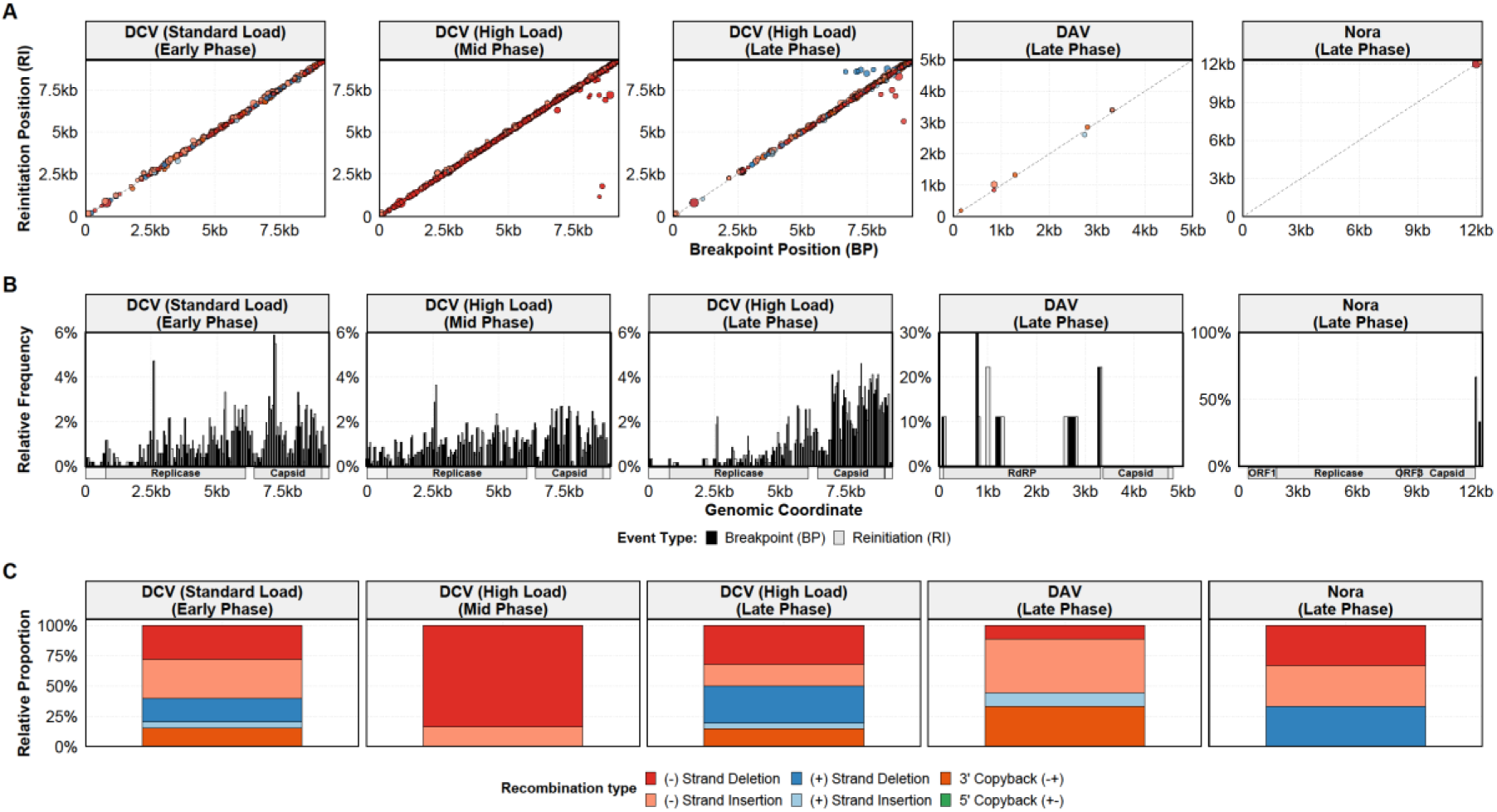
Spatiotemporal dynamics of DVGs during persistent infections. **(A)** Spatiotemporal recombination architecture displaying breakpoint position (BP) versus reinitiation position (RI) across infection stages for DCV (under standard and high load conditions), DAV and Nora virus. Individual points represent unique DVG species, with point sizes scaled by log10 read abundance and colored by recombination type. **(B)** Recombination hotspot density distribution calculated across 100-bp genomic windows, illustrating the relative frequency of breakpoints (black bars) and reinitiation events (gray bars) along the genomic coordinates. **(C)** DVG structural typology distribution showing the relative proportion of recombination types across each viral condition and infection stage.

To determine the spatial distribution of recombination breakpoints and reinitiation events, we mapped their density across the viral genomes using 100-bp sliding windows (Fig. 6B). In DCV, recombination events shifted across infection stages: early and intermediate stages displayed a widely distributed profile characterized by two distinct regional peaks (centering around ∼2.6 kb and ∼7.2–7.8 kb), which progressively concentrated into the capsid gene in the late stage infection. In contrast, DAV recombination events exhibited prominent hotspot peaks distributed along the RdRp gene. For Nora, all identified DVGs were located in the 3’UTR region.

Recombination typology and strand bias varied across infection stages and viral species (Fig. 6C). Restricting our evaluation to the conditions where DVG populations reached detectable thresholds, in standard-load DCV populations at 6 dpe (an additional early-stage time point for DCV), DVGs comprised a mixture of deletions, insertions, and copyback structures. Conversely, in intermediate-stage (12 dpe) high-load DCV populations, negative-sense deletions accounted for the vast majority of DVGs. By the late stage (28 dpe), high-load DCV populations re-established a heterogeneous distribution of deletions, insertions, and copybacks. Late-stage DAV DVGs consisted primarily of negative-sense insertions and 3’ copyback variants, whereas late-stage Nora virus DVGs partitioned evenly among positive-sense deletions, negative-sense deletions, and negative-sense insertions.

### Bloomfield virus maintains a stable genomic architecture and high sequence fidelity

Unlike monopartite ssRNA viruses, segmented dsRNA viruses must coordinate the replication and packaging of multiple distinct genomic molecules while replicating via a double-stranded intermediate. To determine whether persistent infection disrupts segment stoichiometry or drives mutational diversification in a segmented dsRNA virus, we quantified relative segment abundance and sequence variation across nine annotated segments (out of a total of ten, as segment 3 is unavailable) of Bloomfield virus at early (1 dpe), intermediate (12 dpe), and late (36 dpe) infection stages (Figure 7). Segment abundance exhibited a strictly stable, non-equimolar distribution. Normalized to the least abundant segments (S9 and S10), S7 emerged as the most abundant (ratio ∼4–5), followed by S1 and S2 (ratio ∼3), and an intermediate group comprising S4, S5, S6, and S8 (ratio ∼2–2.5). This specific stoichiometric formula remained stable: each segment maintained its proportional abundance across all biological replicates and time points. Alongside this structural constancy, Bloomfield virus maintained robust sequence fidelity (Figure 7B). Across all time points, individual segments consistently harbored fewer than five mutations, and temporal shifts in allele frequency remained tightly constrained below 25%. A singular deviation occurred during the late infection stage, where one biological replicate accumulated a broad swarm of novel variants spanning the entirety of segment S10.

**Fig 7.**
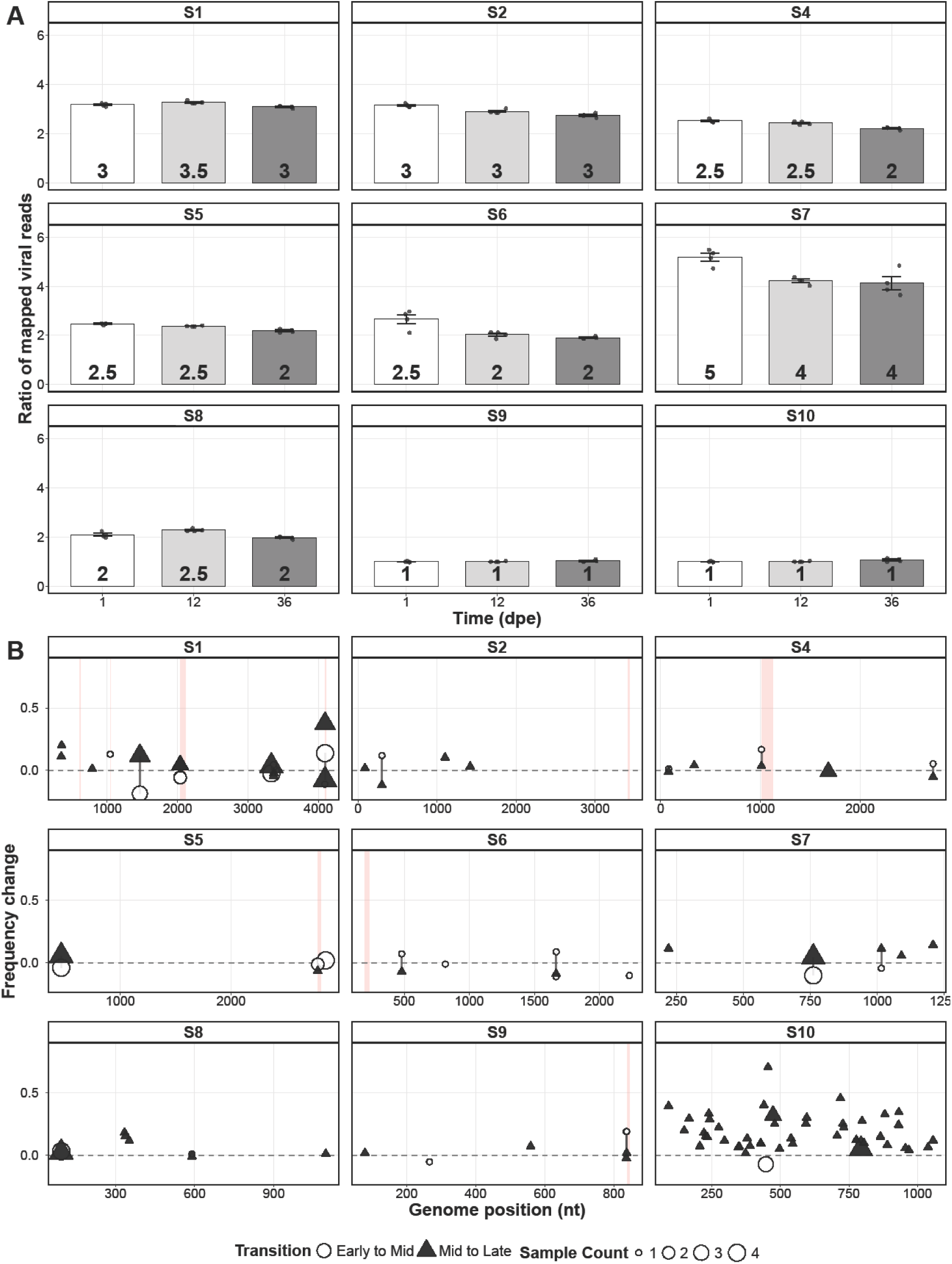
Bloomfield virus maintains a stable genomic formula and high sequence fidelity. **(A)** Relative abundance of the nine analyzed genomic segments of Bloomfield virus at 1, 12, and 36 dpe. Segment abundance is calculated as a read ratio normalized to the baseline of S9 and S10. Bars represent the mean ratio, error bars indicate the standard error of the mean (SEM), and dots denote independent biological replicates. **(B)** Temporal changes in allele frequency (ΔAF) for individual mutations across each genomic segment. White circles denote frequency shifts from early to intermediate infection (1 to 12 dpe); dark triangles indicate shifts from intermediate to late infection (12 to 36 dpe). Symbol size is proportional to the number of biological replicates harboring the specific mutation. Light red vertical bands mask regions of the reference genome containing ambiguous bases (’N’) with insufficient coverage for reliable variant calling.

## DISCUSSION

Long-term viral persistence does not result in a singular evolutionary outcome. By tracking four phylogenetically diverse RNA viruses across the *Drosophila melanogaster* adult lifespan, we demonstrate that persistent infections are highly dynamic systems. Rather than converging on a universal adaptive landscape, each virus follows a distinct evolutionary trajectory dictated by the strict interplay between its specific replication mechanics and the shifting physiological context of the aging host. While these evolutionary dynamics are captured here under standard laboratory conditions, persistent infections occur within a changing environmental context that shapes infection outcomes and influences the emergence and selection of variants in the viral population (Alto et al. 2013, Weger-Lucarelli et al. 2019, Alcaide et al. 2021, González et al. 2021, González et al. 2026b).

The three ssRNA viruses exhibited contrasting modes of adaptation. The evolutionary dynamics of DAV are consistent with clonal interference (Miralles et al. 1999). The continuous generation of a mutant swarm, where high-frequency mutations compete simultaneously on distinct genetic backgrounds, prevents any single variant from reaching fixation. This genetic architecture is a hallmark of large, high-mutation viral populations like HIV and Hepatitis C virus, where the continuous turnover of competing variants facilitates persistence by outpacing host immune recognition (Gerrish & Lenski, 1998; Rambaut et al. 2004, Zanini et al. 2015, Feder et al. 2017, Montoya et al. 2021).

In contrast, DCV evolution under standard-load conditions is governed by strong purifying selection. This genetic homogeneity correlates with a controlled, persistent infection, directly mirroring experimental models where restricted quasispecies diversity attenuates RNA virus pathogenesis (Vignuzzi et al. 2006). However, this equilibrium collapses during high-load infections, triggering mutational diversification. This rapid expansion is analogous to the opportunistic viral diversification observed in immunocompromised human patients, where relaxed immune surveillance removes the biological constraints on viral exploration (Corey et al. 2021).

Nora virus evolution is characterized by delayed positive selection. Following profound evolutionary stasis during early adulthood, the viral population undergoes a late-onset, explosive wave of adaptation driven by large-effect mutations. This delayed diversification is consistent with a rapid viral response to the waning immune competence and physiological decline characteristic of aging *Drosophila* (Zerofsky et al. 2005, Kubiak and Tinsley 2017, Lee et al. 2022). Senescence-dependent expansions are a defining feature of lifelong human infections, such as Cytomegalovirus, which reactivate and adapt by exploiting the declining immune environment of the aging host (Nikolich-Žugich 2008).

Bloomfield virus demonstrates that long-term persistence does not inherently drive a strong diversification of the viral population. While occasional low-level variation occurs, this dsRNA virus preserves a constrained quasispecies structure and an immutable genomic formula across host aging. This constancy is particularly notable because genome formulas are not intrinsically rigid, they can be actively modulated depending on host environments and other factors (Gallet et al. 2022, Bonnamy et al. 2024). The Bloomfield stability confirms that the diverse evolutionary trajectories observed in DAV, DCV, and Nora virus are not driven by the persistent state alone, but by the strict interplay between error-prone ssRNA synthesis, the viral infection strategy, and the shifting physiological status of the host. Ultimately, these results demonstrate that long-term persistence is not a uniform evolutionary state. Instead, viral trajectories diverge sharply based on the viral species, underscoring how specific host– virus combinations structure fitness variance and shape coevolutionary outcomes (Magwire et al. 2012, Belyi et al. 2026).

The generation of defective viral genomes (DVGs) adds an additional structural layer to viral persistence. The spatial restriction of recombination breakpoints (concentrating in the capsid gene of late-stage DCV, the RdRp of DAV, and exclusively within the 3’UTR of Nora virus) indicates that local genomic features, such as secondary RNA structures or polymerase pause sites, govern copy-choice recombination in a species-specific manner (Simon-Loriere & Holmes, 2011). Moreover, DVG abundance and typology scaled with replication magnitude: high-load DCV infections triggered a massive mid-stage expansion of negative-sense deletions before re-establishing structural equilibrium in late infection. The DVG abundance and typology scaled with replication magnitude and exhibited pronounced strand asymmetry parallels recent intra-host observations in nematode RNA viruses (Olmo-Uceda et al., 2026). Notably, 3’ copyback DVGs were present in pathogenic DAV and DCV infections but entirely absent in Nora virus. Because copyback RNAs are potent immunostimulatory ligands (Vignuzzi and López, 2019), their absence in Nora virus aligns with its milder pathogenicity and attenuated host immune activation compared to DAV and DCV (Castelló-Sanjuán et al., 2025). While this study establishes distinct evolutionary trajectories for persistent viruses, certain methodological boundaries contextualize these findings. First, the cross-sectional, destructive sampling required for RNA sequencing impedes tracking the infection trajectory within individual hosts over time. Thus, it remains to be resolved whether the age-induced breakdown of host immunity simply permits opportunistic viral evolution, or if the emergence of high-fitness variants actively drives systemic host senescence. Second, while the sequencing depth successfully captured quasispecies diversity and dominant DVG architectures, ultra-deep sequencing would be required to fully profile the rarest low-frequency recombination events. Finally, the reliance on short-read platforms requires the statistical inference of genetic linkage; future applications of long-read sequencing technologies will allow for the direct physical resolution of full-length viral haplotypes.

In conclusion, our data demonstrate that long-term persistent infections are a dynamic evolutionary playground for viruses, continuously shaped by the interaction between viral exploration of the mutational space, host responses, and host senescence.

## METHODS

### Host and viruses

Fly infection comes from Castelló-Sanjuán et al. (2025). Briefly, *Wolbachia*-free *D. melanogaster* flies were maintained at 25°C on a standard Bloomington cornmeal diet under a 12 h:12 h light cycle. Prior to infection status assessment, flies were confirmed to be free of commonly observed viral contaminants. Experimental groups consisted of stock lines persistently infected with a single RNA virus (DAV, DCV, Nora virus, or Bloomfield virus) established in Nigg et al. (2024). Sample-matched intra-host viral loads were retrieved from Castelló-Sanjuán et al. (2025) to classify standard-versus high-load DCV infections and to evaluate correlations with cumulative mutational burden

### Viral genomes and annotations

Reference viral genomes utilized for all alignments and structural annotations were retrieved from the NCBI GenBank database. For the positive-sense single-stranded RNA (ssRNA) viruses, the references were: *Drosophila A virus* isolate HD (GenBank: FJ150422.1), *Drosophila C virus* isolate SMN (GenBank: OK188767.1), and Nora virus isolate SMN (GenBank: OK188768.1). For the multi-segmented double-stranded RNA (dsRNA) Bloomfield virus, nine genomic segments were analyzed corresponding to GenBank accessions KP714090.1 (Segment 1), KP714091.1 (Segment 2), and KP714092.1 through KP714098.1 (Segments 4 to 10, respectively).

Genomic annotations defining untranslated regions (UTRs) and coding sequences (CDS) were strictly mapped according to their respective GFF3 profiles. For DAV (4,806 nt), structural constraints were mapped across the RNA-dependent RNA polymerase (RdRp, nt 88–3315) and the structural protein precursor (nt 3368–4696). For DCV (9,264 nt), annotations encompassed ORF-1 encoding the replicase polyprotein (nt 799–6078) and ORF-2 encoding the capsid polyprotein (nt 6267–8972). For Nora virus (12,333 nt), specific structural mappings were applied to ORF1 (nt 479–1906), the replicase polyprotein (ORF2, nt 1900–8214; harboring the Helicase domain at nt 2962–3772 and RdRp domain at nt 7183– 8098), ORF3 (nt 8189–9058), and the main structural capsid protein (ORF4, nt 9144–11939).

### High-throughput sequencing

All genomic analyses of standard-load infections were performed using the longitudinal RNA-seq datasets described in Castelló-Sanjuán et al. (2025). Briefly, four biological replicates per treatment group were prepared, each consisting of pooled RNA from three to five individual flies (raw reads available under BioProject PRJNA1235228). For the newly generated high-load DCV dataset, biological replicates were constructed by pooling RNA from two individual flies exhibiting equivalently high viral loads. These high-load libraries were sequenced using an identical Illumina NextSeq 500 platform pipeline. Raw sequence data for the high-load samples have been deposited in the NCBI Sequence Read Archive under BioProject PRJNA1527392.

### Processing of sequencing reads

Raw sequencing reads were quality-filtered and trimmed using Cutadapt (Martin 2011) to remove adapters and technical artifacts. To ensure high-quality alignments, reads shorter than 40 nucleotides were discarded, and 17 bases were strictly trimmed from the 5’ end. Trimmed reads were aligned to their respective viral reference genomes (DAV, DCV, Nora virus, and Bloomfield virus) using HISAT2 (Kim et al. 2019) with spliced alignments disabled. Alignments were processed, sorted, and indexed using SAMtools (Li et al., 2009). High-confidence mutations were called using LoFreq (Wilm et al. 2012), and per-position base compositions were extracted using pysamstats. To strictly isolate biological variation from sequencing noise, sub-consensus variants were filtered using bcftools to retain only those with an allele frequency (AF) ≥ 0.01. Defective viral genomes (DVGs) were identified, quantified, and structurally classified directly from the sorted alignment files using DVGfinder (Olmo-Uceda et al., 2022).

### Nucleotide diversity

Nucleotide diversity (*π*) was quantified across the viral genomes from aligned sequencing reads using per-nucleotide depth matrices generated via pysamstats. Polymorphic sites were defined by a dual threshold: a minimum sequencing depth of 30 × and a minor allele frequency ≥ 0.01. For each validated site *i*, site-specific nucleotide diversity (*π_i_*) was calculated using the unbiased finite-sample estimator across all four nucleotide bases:

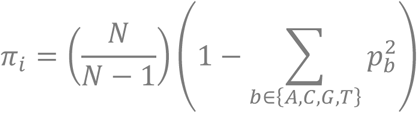

where *N* represents total sequencing depth at position *i*, and *p_b_* denotes the relative frequency of base *b* ∈ {*A*, *C*, *G*, *T*}. The resulting positional *π_i_* values were averaged across defined genomic regions (assigning a value of zero to monomorphic or sub-threshold sites) to map the spatial distribution of mutational constraints.

### Estimation of mutational fitness effects and genetic linkage

To identify parallel adaptive mutations and quantify their intra-host fitness impacts, we filtered for variants detected in at least three biological replicates at a minimum of one infection time point. Allele frequencies (*AF*, scaled from 0 to 1) were bounded at 0.005 and 0.995 to accommodate boundary values and transformed using a base-10 logit function:

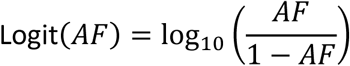

For each variant with at least six longitudinal observations, temporal trajectories were modeled using linear mixed-effects models implemented in the R packages *lme4* and *lmerTest* (Bates et al. 2015, Kuznetsova et al. 2017), fitting Logit(*AF*) as a function of time (days post-eclosion) as a fixed effect and biological replicate as a random intercept (Logit(*AF*) ∼ Time + (1|Replicate)) via maximum likelihood (falling back to ordinary least-squares regression when random-effect variance approached zero). Relative fitness effects per day were derived by exponentiating the fixed-effect slope (*w* = 10*^β^*^Time^ ) and expressed either as relative fitness compared to wild-type (*w*or as log_2_-transformed fitness effects (log_2_(*w*)). Statistical significance of the temporal slope was evaluated using Satterthwaite’s degrees of freedom method and adjusted for multiple testing via the Benjamini–Hochberg false discovery rate (FDR).

To infer genetic linkage and clonal interference across the viral quasispecies, parallel variants present in at least three samples with an allele frequency standard deviation > 0.01 were retained. Pairwise Pearson correlation coefficients (*r*) between untransformed allele frequencies across all biological replicates and time points were computed using the *Hmisc* package in R.

### Protein structure and thermodynamic modeling

To evaluate the structural topography of the identified adaptive mutations, we utilized the 2.7 Å cryo-EM structure of the Nora virus capsid (Laurinmäki et al., 2020). The 3D structures of the remaining viral RNA-dependent RNA polymerases (RdRps) and capsid proteins were modeled using AlphaFold3 (Abramson et al. 2024), with OpenFold3 (Ahdritz et al. 2024) specifically deployed for the Nora virus replicase. Predicted structures were filtered using a combination of global (pTM) and local (pLDDT) confidence metrics. Full-length models with low global confidence (pTM < 0.70) were retained only if core structural regions scored pLDDT > 90, strictly limiting subsequent analyses to high-confidence domains. We calculated thermodynamic stability changes (ΔΔG) induced by single amino acid substitutions using ThermoMPNN (Dieckhaus et al., 2024). Structural visualization and annotation were performed in ChimeraX 1.9 (Meng et al., 2023).

### Statistical analysis

All statistical analyses and data visualizations were executed in R (version 4.3.2) within the RStudio environment (version 2026.8.0.187), utilizing the ggplot2 package for graphics. The specific statistical tests deployed for each analysis, along with their corresponding sample sizes are defined in the figure legends. Statistical significance was established at *P* < 0.05 for all analyses and symbols denote * P < 0.05, ** P < 0.01, *** P < 0.001.

## DATA AVAILABILITY

RNA sequencing data analyzed in this study are deposited in the NCBI Sequence Read Archive under BioProject accessions PRJNA1235228 (standard-load infection data from Castelló-Sanjuán et al., 2025) and PRJNA1527392 (high-load DCV samples sequenced for this study). All other data supporting the findings of this study are available within the paper and its Supplementary Material (Supplementary Files 1–4). Custom R scripts for data analysis and figure generation are available from the corresponding authors upon reasonable request.

## ACKNOWLEDGMENTS

We thank all members of the Saleh laboratory for helpful discussions and constructive feedback on this work. We acknowledge the use of Gemini (Google) for assistance with script refinement, manuscript editing, and proofreading.

## AUTHOR CONTRIBUTIONS

Conceptualization: R.G. and M-C.S. Methodology: R.G. and A.B. Investigation: R.G., A.B., H.B., M.C-S. Visualization: A.B. and R.G. Supervision: M-C.S. and R.G. Writing—original draft: R.G. Writing—review and editing: R.G., M.C-S., A.B. Funding acquisition: M-C.S. Data curation: R.G. Project administration: R.G. and M-C.S. Formal analysis: R.G. and A.B.

## SUPPLEMENTARY FIGURES

**Supplementary Fig 1.**
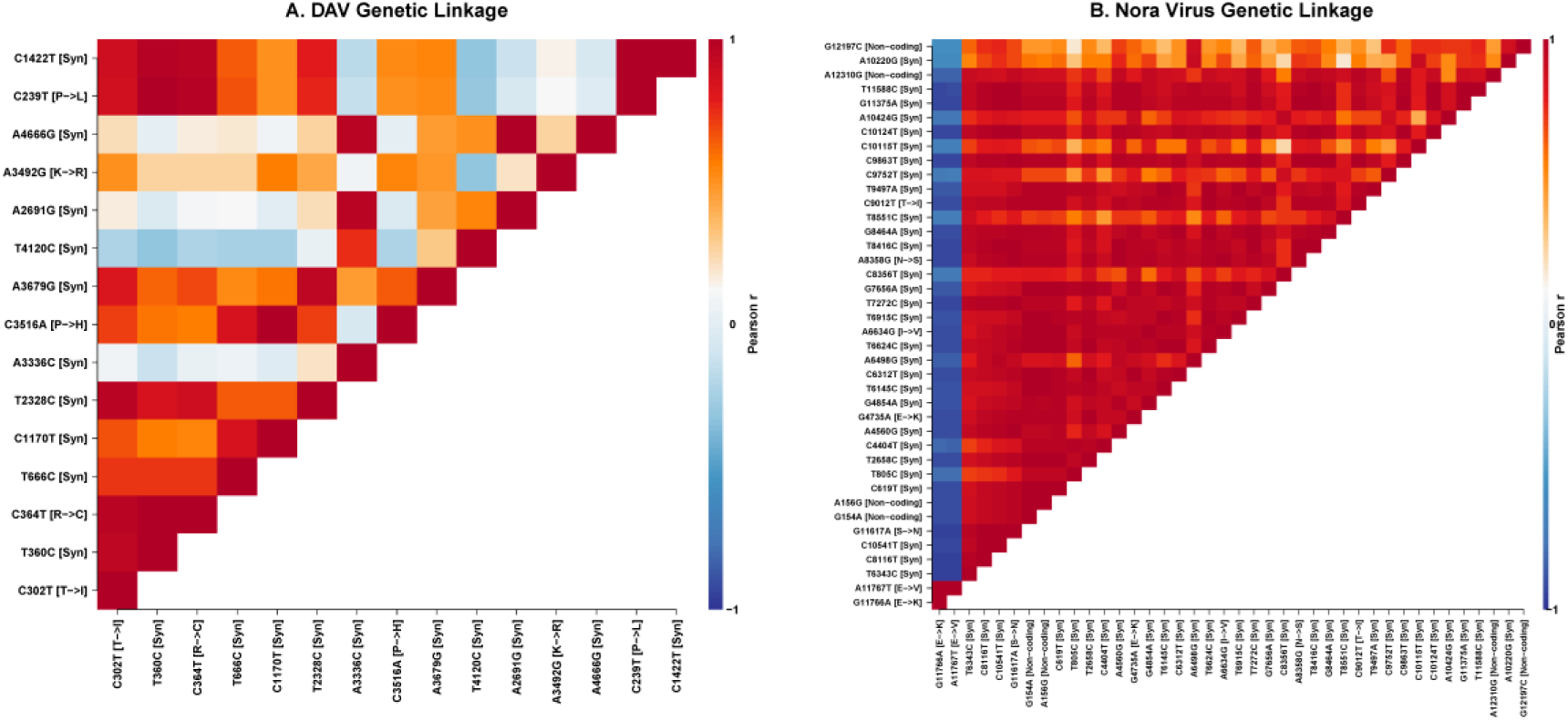
Genetic linkage matrices of adaptive mutations. Correlation structure among significant adaptive mutations (detected in ≥3 samples with SD > 0.01) identified during persistent infections with DAV (panel A) and Nora virus (panel B). Color scale indicates Pearson’s correlation coefficient (r), ranging from strong positive correlation (red, r = 1) to strong negative correlation (blue, r = -1)

**Supplementary Fig 2.**
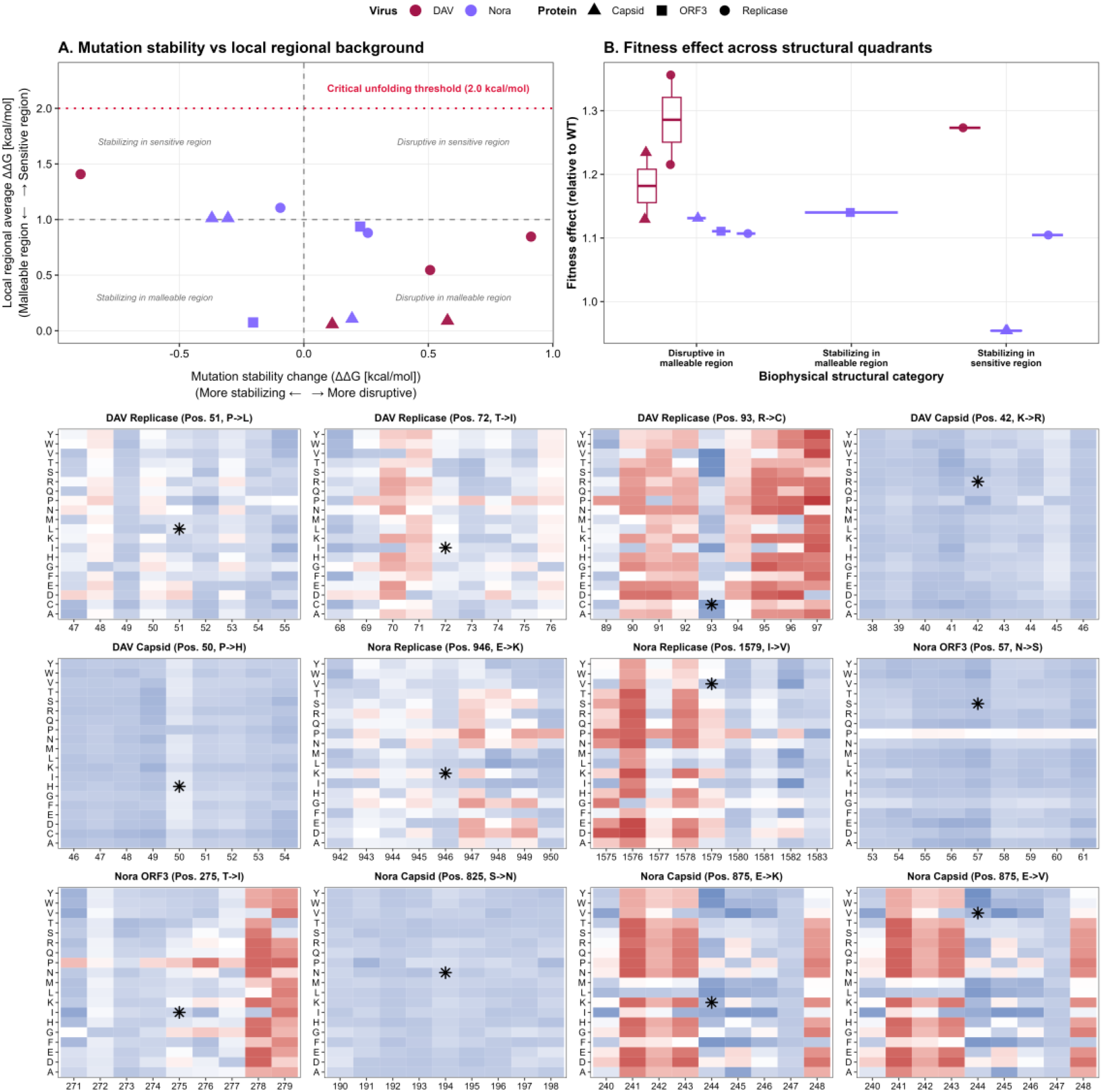
Biophysical stability landscapes, structural quadrants, and comprehensive mutational scanning heatmaps for DAV and Nora virus proteins. **(A)** Mutation stability change (ΔΔG, kcal/mol) plotted against the local regional background (malleable versus sensitive regions), bounded by the critical unfolding threshold of 2.0 kcal/mol. **(B)** Fitness effects relative to wild-type (WT) across defined biophysical structural categories (disruptive in malleable/sensitive regions and stabilizing in malleable/sensitive regions). (Lower Panels) Positional saturation mutagenesis and thermodynamic stability matrices for specific residues across DAV (Replicase and Capsid) and Nora virus (Replicase, Capsid, and ORF3) proteins. Heatmaps display amino acid substitutions (single-letter codes on the y-axis against residue positions of the peptide on the x-axis), indicating stabilization (blue shades) versus destabilization/disruption (red shades), with wild-type residues denoted by asterisks (*).

**Supplementary Fig 3.**
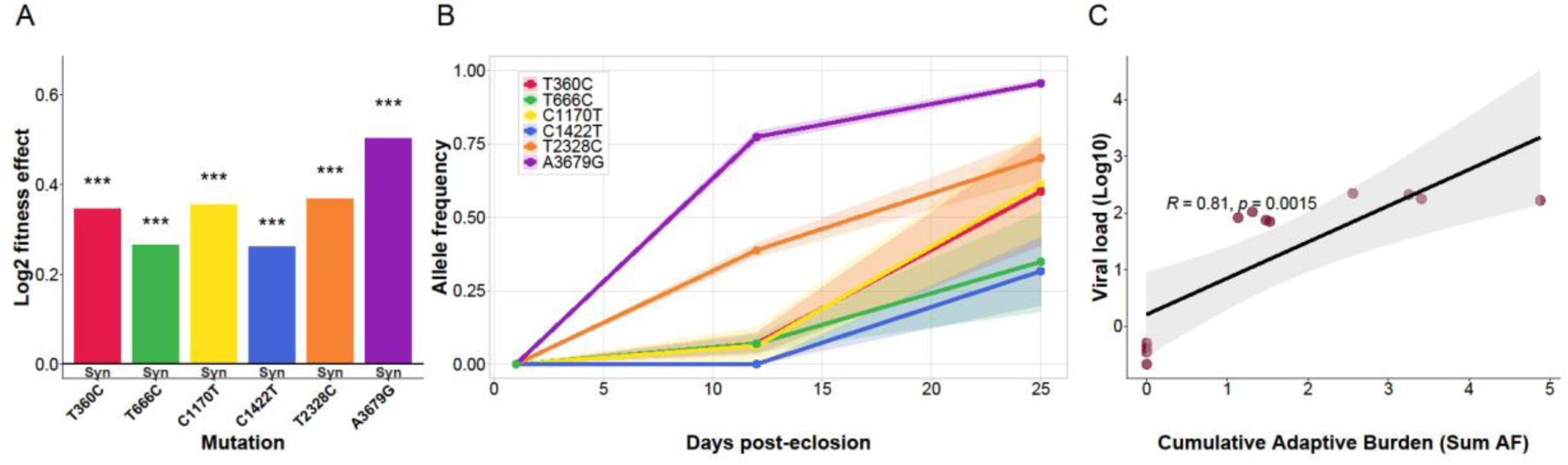
Fitness impacts of adaptive synonymous mutations in DAV. **(A)** Fitness effects of all 6 significant synonymous adaptive mutations during DAV persistence. Asterisks denote statistical significance. **(B)** Temporal allele frequency trajectories of these variants across days post-eclosion. Shaded regions represent the standard error of the mean across independent biological replicates. **(C)** Pearson correlation between the cumulative adaptive mutational burden (sum of allele frequencies) and intra-host viral load. The solid black line indicates the linear regression fit, and the shaded area represents the 95% confidence interval (R=0.81, p=0.0015).

**Supplementary Fig 4.**
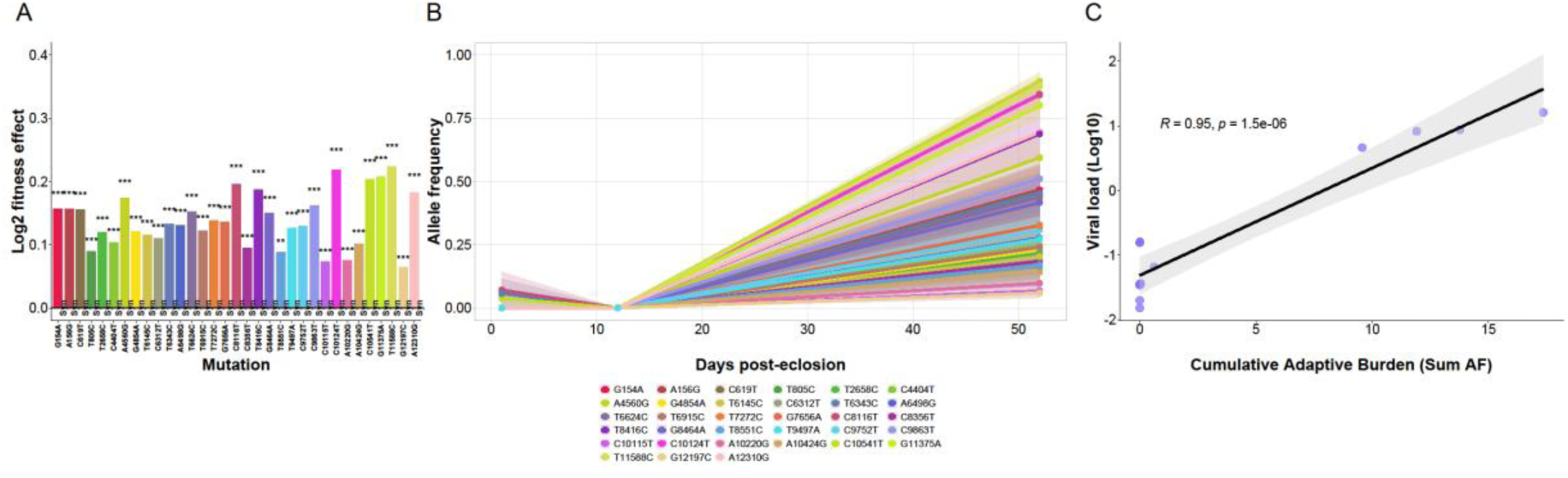
Fitness impacts of adaptive synonymous mutations in Nora virus. **(A)** Fitness effects of all 33 significant synonymous and non-coding adaptive mutations during Nora persistence. Asterisks denote statistical significance. **(B)** Temporal allele frequency trajectories of these variants across days post-eclosion. Shaded regions represent the standard error of the mean across independent biological replicates. **(C)** Pearson correlation between the cumulative adaptive mutational burden (sum of allele frequencies) and intra-host viral load. The solid black line indicates the linear regression fit, and the shaded area represents the 95% confidence interval (R=0.95, p < 0.0001).

## SUPPLEMENTARY FILES

**Supplementary File 1.** Dataset of all genomic variants identified across persistent infection time points for Drosophila A virus, Drosophila C virus, and Nora virus.

**Supplementary File 2.** Dataset of all significant mutations and their synonymous or non-synonymous changes for Drosophila A virus and Nora virus.

**Supplementary File 3.** Dataset of all high-confidence genomic variants for high-versus standard-load Drosophila C virus (DCV) populations at 6, 12, and 28 dpe.

**Supplementary File 4.** Dataset of all DVGs detected.

## REFERENCES

1. Abramson J, Adler J, Dunger J et al. Accurate structure prediction of biomolecular interactions with AlphaFold 3. Nature 2024;630(8016):493–500. 10.1038/s41586-024-07487-w.

2. Ahdritz G, Bouatta N, Floristean C et al. OpenFold: retraining AlphaFold2 yields new insights into its learning mechanisms and capacity for generalization. Nat Methods 2024;21(8):1514–24. 10.1038/s41592-024-02272-z.

3. Alcaide C, Sardanyés J, Elena SF et al. Increasing temperature alters the within-host competition of viral strains and influences virus genetic variability. Virus Evolution 2021;7(1):veab017. 10.1093/ve/veab017.

4. Alto BW, Wasik BR, Morales NM et al. Stochastic temperatures impede rna virus adaptation: temperature adaptation of an RNA virus. Evolution 2013;67(4):969–79. 10.1111/evo.12034.

5. Bates D, Mächler M, Bolker B et al. Fitting Linear Mixed-Effects Models Using lme4. J Stat Soft 2015;67(1). 10.18637/jss.v067.i01.

6. Belyi A, Du Y, Wilson AJ et al. Host–virus interactions dominate fitness and conceal additive genetic variation. PLoS Biol 2026;24(10):e3003836. 10.1371/journal.pbio.3003836.

7. Bonnamy M, Brousse A, Pirolles E et al. The genome formula of a multipartite virus is regulated both at the individual segment and the segment group levels. PLoS Pathog 2024;20(1):e1011973. 10.1371/journal.ppat.1011973.

8. Bonneaud C, Longdon B. Emerging pathogen evolution: Using evolutionary theory to understand the fate of novel infectious pathogens. EMBO Rep 2020;21(9):EMBR202051374. 10.15252/embr.202051374.

9. Castelló-Sanjuán M, González R, Romoli O et al. Persistent viral infections impact key biological traits in Drosophila melanogaster. PLoS Biol 2025;23(10):e3003437. 10.1371/journal.pbio.3003437.

10. Castiglioni VG, Olmo-Uceda MJ, Martín S et al. Experimental evolution of an RNA virus in Caenorhabditis elegans. *Infection*, Genetics and Evolution 2024;123:105623. 10.1016/j.meegid.2024.105623.

11. Corey L, Beyrer C, Cohen MS et al. SARS-CoV-2 Variants in Patients with Immunosuppression. N Engl J Med 2021;385(6):562–6. 10.1056/NEJMsb2104756.

12. Dieckhaus H, Brocidiacono M, Randolph NZ et al. Transfer learning to leverage larger datasets for improved prediction of protein stability changes. Proc Natl Acad Sci USA 2024;121(6):e2314853121. 10.1073/pnas.2314853121.

13. Dionne MS, Schneider DS. Models of infectious diseases in the fruit fly *Drosophila melanogaster*. Disease Models & Mechanisms 2008;1(1):43–9. 10.1242/dmm.000307.

14. Domingo E. Long-term virus evolution in nature. In: Virus as Populations. n.p.: Elsevier, 2020, 225–61. 10.1016/B978-0-12-816331-3.00007-6.

15. Farci P, Shimoda A, Coiana A et al. The Outcome of Acute Hepatitis C Predicted by the Evolution of the Viral Quasispecies. Science 2000;288(5464):339–44. 10.1126/science.288.5464.339.

16. Feder AF, Kline C, Polacino P et al. A spatio-temporal assessment of simian/human immunodeficiency virus (SHIV) evolution reveals a highly dynamic process within the host. PLoS Pathog 2017;13(5):e1006358. 10.1371/journal.ppat.1006358.

17. Frielle DW, Huang DD, Youngner JS. Persistent infection with influenza A virus: Evolution of virus mutants. Virology 1984;138(1):103–17. 10.1016/0042-6822(84)90151-X.

18. Gallet R, Di Mattia J, Ravel S et al. Gene copy number variations at the within-host population level modulate gene expression in a multipartite virus. Virus Evolution 2022;8(2):veac058. 10.1093/ve/veac058.

19. Gerrish PJ, Lenski RE. The fate of competing beneficial mutations in an asexual population. Genetica 1998;102–103(1–6):127–44.

20. Goic B, Saleh MC. Living with the enemy: viral persistent infections from a friendly viewpoint. Curr Opin Microbiol 2012;15(4):531–7. 10.1016/j.mib.2012.06.002.

21. González R, Butković A, Escaray FJ et al. Plant virus evolution under strong drought conditions results in a transition from parasitism to mutualism. Proc Natl Acad Sci USA 2021;118(6):e2020990118. 10.1073/pnas.2020990118

22. González R, Castelló-Sanjuán M, Saleh MC. Enteric viral infections promote systemic accelerated aging in Drosophila. Sci Adv 2026a;12(23):eaec1735. 10.1126/sciadv.aec1735.

23. González R, Castelló-Sanjuán M, Butković A et al. Parasitism to mutualism shift during a viral infection in Drosophila under thermal stress. bioRxiv 2026b 10.64898/2026.10.31.763142.

24. Holland JJ, Grabau EA, Jones CL et al. Evolution of multiple genome mutations during long-term persistent infection by vesicular stomatitis virus. Cell 1979;16(3):495–504. 10.1016/0092-8674(79)90024-2.

25. Holmes EC. The Evolution and Emergence of RNA Viruses. n.p.: Oxford University Press Oxford, 2009. 10.1093/oso/9780199211128.001.0001.

26. Kim D, Paggi JM, Park C et al. Graph-based genome alignment and genotyping with HISAT2 and HISAT-genotype. Nat Biotechnol 2019;37(8):907–15. 10.1038/s41587-019-0201-4.

27. Kubiak M, Tinsley MC. Sex-Specific Routes To Immune Senescence In Drosophila melanogaster. Sci Rep 2017;7(1):10417. 10.1038/s41598-017-11021-6.

28. Kuznetsova A, Brockhoff PB, Christensen RHB. lmerTest Package: Tests in Linear Mixed Effects Models. J Stat Soft 2017;82(13). 10.18637/jss.v082.i13.

29. Lauring AS, Andino R. Quasispecies Theory and the Behavior of RNA Viruses. PLoS Pathog 2010;6(7):e1001005. 10.1371/journal.ppat.1001005.

30. Laurinmäki P, Shakeel S, Ekström JO et al. Structure of Nora virus at 2.7 Å resolution and implications for receptor binding, capsid stability and taxonomy. Sci Rep 2020;10(1):19675. 10.1038/s41598-020-76613-1.

31. Lee KA, Flores RR, Jang IH et al. Immune Senescence, Immunosenescence and Aging. Front Aging 2022;3:900028. 10.3389/fragi.2022.900028.

32. Lezcano OM, Fuhrmann L, Ramakrishnan G et al. Parallel evolution and enhanced virulence upon in vivo passage of an RNA virus in *Drosophila melanogaster*. Virus Evolution 2023;9(2):vead074. 10.1093/ve/vead074.

33. Li H, Handsaker B, Wysoker A et al. The Sequence Alignment/Map format and SAMtools. Bioinformatics 2009;25(16):2078–9. 10.1093/bioinformatics/btp352.

34. Magwire MM, Fabian DK, Schweyen H et al. Genome-Wide Association Studies Reveal a Simple Genetic Basis of Resistance to Naturally Coevolving Viruses in Drosophila melanogaster. PLoS Genet 2012;8(11):e1003057. 10.1371/journal.pgen.1003057.

35. Martin M. Cutadapt removes adapter sequences from high-throughput sequencing reads. EMBnet j 2011;17(1):10. 10.14806/ej.17.1.200.

36. Meng EC, Goddard TD, Pettersen EF et al. UCSF ChimeraX : Tools for structure building and analysis. Protein Science 2023;32(11):e4792. 10.1002/pro.4792.

37. Miralles R, Gerrish PJ, Moya A et al. Clonal Interference and the Evolution of RNA Viruses. Science 1999;285(5434):1745–7. 10.1126/science.285.5434.1745.

38. Mongelli V, Lequime S, Kousathanas A et al. Innate immune pathways act synergistically to constrain RNA virus evolution in *Drosophila melanogaster*. Nat Ecol Evol 2022;6(5):565–78. 10.1038/s41559-022-01697-z.

39. Montoya V, Howe AYM, Dong WY et al. Intra-host evolutionary dynamics of the hepatitis C virus among people who inject drugs. Sci Rep 2021;11(1):9986. 10.1038/s41598-021-88132-8.

40. Nigg JC, Castelló-Sanjuán M, Blanc H et al. Viral infection disrupts intestinal homeostasis via Sting-dependent NF-κB signaling in Drosophila. Current Biology 2024;34(13):2785–2800.e7. 10.1016/j.cub.2024.05.009.

41. Nikolich-Žugich J. Ageing and life-long maintenance of T-cell subsets in the face of latent persistent infections. Nat Rev Immunol 2008;8(7):512–22. 10.1038/nri2318.

42. Obbard DJ, Shi M, Roberts KE et al. A new lineage of segmented RNA viruses infecting animals. Virus Evol 2020;6(1):vez061. 10.1093/ve/vez061.

43. Olmo-Uceda MJ, Muñoz-Sánchez JC, Lasso-Giraldo W et al. DVGfinder: A Metasearch Tool for Identifying Defective Viral Genomes in RNA-Seq Data. Viruses 2022;14(5):1114. 10.3390/v14051114.

44. Olmo-Uceda MJ, Herek D, Castiglioni VG et al. The hidden diversity: Replication-strand asymmetry shapes short-term nucleotide and structural variation in an RNA virus. bioRxiv 2026. 10.64898/2026.09.04.749430.

45. Rambaut A, Posada D, Crandall KA et al. The causes and consequences of HIV evolution. Nat Rev Genet 2004;5(1):52–61. 10.1038/nrg1246.

46. Rezelj VV, Carrau L, Merwaiss F et al. Defective viral genomes as therapeutic interfering particles against flavivirus infection in mammalian and mosquito hosts. Nat Commun 2021;12(1):2290. 10.1038/s41467-021-22341-7.

47. Simon-Loriere E, Holmes EC. Why do RNA viruses recombine? Nat Rev Microbiol 2011;9(8):617–26. 10.1038/nrmicro2614.

48. Vignuzzi M, López CB. Defective viral genomes are key drivers of the virus–host interaction. Nat Microbiol 2019;4(7):1075–87. 10.1038/s41564-019-0465-y.

49. Vignuzzi M, Stone JK, Arnold JJ et al. Quasispecies diversity determines pathogenesis through cooperative interactions in a viral population. Nature 2006;439(7074):344–8. 10.1038/nature04388.

50. Wallace MA, Coffman KA, Gilbert C et al. The discovery, distribution, and diversity of DNA viruses associated with Drosophila melanogaster in Europe. Virus Evol 2021;7(1):veab031. 10.1093/ve/veab031.

51. Wallace MA, Obbard DJ. Naturally occurring viruses of Drosophila reduce offspring number and lifespan. Proc Biol Sci 2024;291(2023):20240518. 10.1098/rspb.2024.0518.

52. Webster CL, Longdon B, Lewis SH et al. Twenty-Five New Viruses Associated with the Drosophilidae (Diptera). Evol Bioinform Online 2016;12(Suppl 2):13–25. 10.4137/EBO.S39454.

53. Webster CL, Waldron FM, Robertson S et al. The Discovery, Distribution, and Evolution of Viruses Associated with Drosophila melanogaster. PLoS Biol 2015;13(7):e1002210. 10.1371/journal.pbio.1002210.

54. Weger-Lucarelli J, Carrau L, Levi LI et al. Host nutritional status affects alphavirus virulence, transmission, and evolution. PLoS Pathog 2019;15(11):e1008089. 10.1371/journal.ppat.1008089.

55. Wilm A, Aw PPK, Bertrand D et al. LoFreq: a sequence-quality aware, ultra-sensitive variant caller for uncovering cell-population heterogeneity from high-throughput sequencing datasets. Nucleic Acids Research 2012;40(22):11189–201. 10.1093/nar/gks918.

56. Zanini F, Brodin J, Thebo L et al. Population genomics of intrapatient HIV-1 evolution. eLife 2015;4:e11282. 10.7554/eLife.11282.

57. Zerofsky M, Harel E, Silverman N et al. Aging of the innate immune response in *Drosophila melanogaster*. Aging Cell 2005;4(2):103–8. 10.1111/j.1474-9728.2005.00147.x.

